# The Naïve Bayes Classifier++ as an Out-of-Distribution Detector of Novel Taxa

**DOI:** 10.1101/2025.10.18.683227

**Authors:** Keyush Ramjattun, Gail L. Rosen

## Abstract

Detecting sequences from novel taxa remains a key challenge in metagenomic classification, as reference databases rarely capture the full extent of microbial diversity. We investigate the Naïve Bayes Classifier++ (NBC++) as an out-of-distribution (OOD) detector by analyzing its log-likelihood scores across simulated and real metagenomic datasets. By partitioning reference databases and introducing taxonomic novelty, we derive thresholds that distinguish known from unknown reads at multiple taxonomic levels. These thresholds remain consistent across database sizes, indicating that once a lineage is represented, novelty detection performance stabilizes. Applied to a human gut metagenome, the thresholds reflect differences in database density and classification confidence. This work characterizes how NBC++ responds to novelty and illustrates its use in evaluating unclassified metagenomic reads.

## Introduction

Metagenomics has transformed our ability to profile microbial communities, yet one of the biggest challenges is detecting sequences originating from novel taxa—organisms absent from existing reference databases. It remains a difficult problem due the challenge of identifying something unknown from a reference point of your known. Compounding that, is the taxonomic hierarchy of kingdoms, where each clade may be evolving at a different evolutionary rate. In machine learning, this problem is known as “out-of-distribution” (OOD) detection Yang et al. [2024], Tran [2025]. Large-scale surveys consistently reveal that a meaningful fraction of metagenomic reads cannot be classified. Modha et al. [2022] systematically analyzed human metagenomes and found that reads, which appeared across multiple studies, resulted in”unknown contigs” that assembled into novel viral genomes that bear no similarity to sequences in the public databases. Similar evidence is reported in global-scale metagenomic studies, which highlight extensive “functional dark matter” in the form of uncharacterized protein families Pavlopoulos et al. [2023]. These findings show that unclassified sequences represent real biological diversity rather than random sequencing artefacts.

Metagenomic taxonomic tools have tried to mitigate novelty detection. In Kraken2 Wood et al. [2019], for example, any query sequence that does not contain “enough k-mers” (a threshold that can be defined) belonging to a taxa in the reference database cannot be classified and is therefore marked as “unclassified”. Choosing the appropriate confidence threshold is a trade-off between increasing accuracy vs. the proportion of classified reads that are lost. Also, the percent of reads that could be classified increases as a function of the database size Wright et al. [2023], Liu et al. [2024]. When NBC incorporates minimizers like Kraken2, it is able to obtain a reasonable confidence threshold to filter unknown reads Lu et al. [2024]. However, “unknown reads” were only simulated from the *Arabidopsis thaliana* plant, which may limit the feasibility on unknown microbial sequences.

Therefore, a natural problem to benchmark metagenomic classifiers is generating feasible datasets that take into consideration various types of novelty. Nickols et.al test over different types of novelty: a) taxa missing from a database, newly added NCBI taxa since the last database update, and completely novel species genome bins (SGBs) without NCBI IDs. Taxonomic classifier methods performed better on higher levels of taxonomy than lower levels, especially for real samples Nickols et.al. One of the Nickols et al.’s suggestions is to use MAGs in reference databases, which is also supported by another study Smith et al. [2022]. As more genomes are added to a database, they do not necessarily represent more diversity. Therefore, it is important to have a way to measure diversity growth within clades and pangenomes. A delta measure, based on compression, has been introduced and extended to suggest optimal k-mers, which can be critical to minimizer hash-based methods Bonnie et al. [2024].

In this study, we examine how well the metagenomic NBC++ can identify novel taxa. NBC is competitive and especially, for 16S, newer deep learning architectures cannot outperform it Ziemski et al. [2021]. While there are other classifiers, we want to experiment with NBC++’s score, a log-likelihood probability, that can be used to intelligently compare the probability of sequence reads against a database, and see how well it extends to novelty detection of various taxa. This will be vital to NBC++, since it is an incremental-class learner van de Ven et al. [2022] and to incrementally and learn to distinguish between a growing number of objects or classes will be vital, especially with novel classes being identified before being integrated.

### Methodology

Our goal was to derive a log-likelihood threshold that can be used to identify novel taxa not present in a reference database. The general workflow involved partitioning a database, training NBC, classifying a test set, and calculating a threshold using ROC analysis. This process was applied independently to a “basic” database containing one genome per species and a much larger “extended” database with multiple genomes per species, each requiring specific variations during the data preparation stage.

#### Initial Database Filtering

Both databases were first prepared at the phylum, class, order, and family level. To ensure adequate representation, classes with a low number of genomes were removed. The minimum threshold was set to 30 representatives for the “basic” database and 400 for the “extended” database.

#### Training and Holdout Set Partitioning

After the initial filtering, the remaining genomes were partitioned to create “known” (training) and “unknown” (holdout) sets for the subsequent ROC analysis. The partitioning method differed between the two databases to account for their distinct structures. For the basic database, which contains only one genome per species, a simple random split was performed where 50% of the species were assigned to the training set and the other 50% formed the holdout set. In contrast, the extended database, with multiple genomes per species, required a different approach to ensure the training set was balanced. For these experiments, 50% of the taxonomic classes were first randomly selected, and then 400 genomes were randomly sampled from within each of those selected classes. All genomes from the unselected classes were subsequently designated as the holdout set. This entire partitioning process was repeated five times for each database to generate independent trials for statistical robustness.

#### Model Training and Classification

For each trial, NBC++ was trained on the corresponding training set for various k-mer lengths (3, 6, 9, 12, 15). These models were then used to classify a standardized test set containing reads from both the training (“known”) and holdout (“unknown”) sets, generating a log-probability score for each read.

To derive a novelty threshold, we scored simulated reads generated from all reference species (1 genome for Basic and many genomes per species for Extended) as in Duan et al. [2024]. Therefore, a proportion of the test reads were from species in the training database, and many were from species out-of-distribution. The InSilicoSeq simulator Gourlé et al. [2018] generated 100 reads per species. Then, each simulated read was classified using the trained models to obtain a log-probability score – the log-probability of that read given each training species. Then the read was assigned the maximum likelihood probability as the final score. If the read was computed on a training database that contained the species belonging to the same taxa as the read, its max log-probability score was put into a “known bin”. Conversely, if the read’s taxa was not in the training database, the maximum likelihood score was assigned to an “unknown bin”.

#### ROC analysis

ROC analysis was then performed to find the optimal threshold to separate the “known” reads, that are related to the training data, from the “unknown” reads, from the holdout set. The optimal log-probablity threshold was then determined as the point on the ROC curves that maximized Youden’s J statistic.

#### Real-world validation

To validate our method, we analyzed a human gut metagenomic dataset (SRA ID: SRS105153) described in Zhao et al. [2020] and Nasko et al. [2018]. In contrast to the fixed-length sequences used for model training (126 bp), the real-world sample consists of reads with varying lengths. To address this, we first apply a length-normalization to the novelty threshold for each read. We then classifed each normalized read using the 9-mer models derived from both the basic and extended databases. This process allowed us to assess the practical performance of our novelty detection thresholds and directly compare the models’ outputs on complex, real-world data.

## Results and Discussion

We evaluate the performance of our novelty detection method across a range of key parameters, specifically investigating the impact of k-mer length and database scale. The results of this evaluation are then used to assess our framework’s practical application on the human gut metagenome described in Zhao et al. [2020] and Nasko et al. [2018].

### K-mer length and classification accuracy

We assessed NBC’s ability to distinguish “known” sequences from holdout “unknown” sequences. A noticeable trend emerged for both the basic and extended databases: as the k-mer length increased from 3 to 15, the classifier’s performance improved significantly. There is a consistent increase in the Area Under Curve (AUC) values from the ROC analysis, confirming that longer, more specific k-mers provide greater distinguishing power for classification (Fig 1).

**Figure 1.**
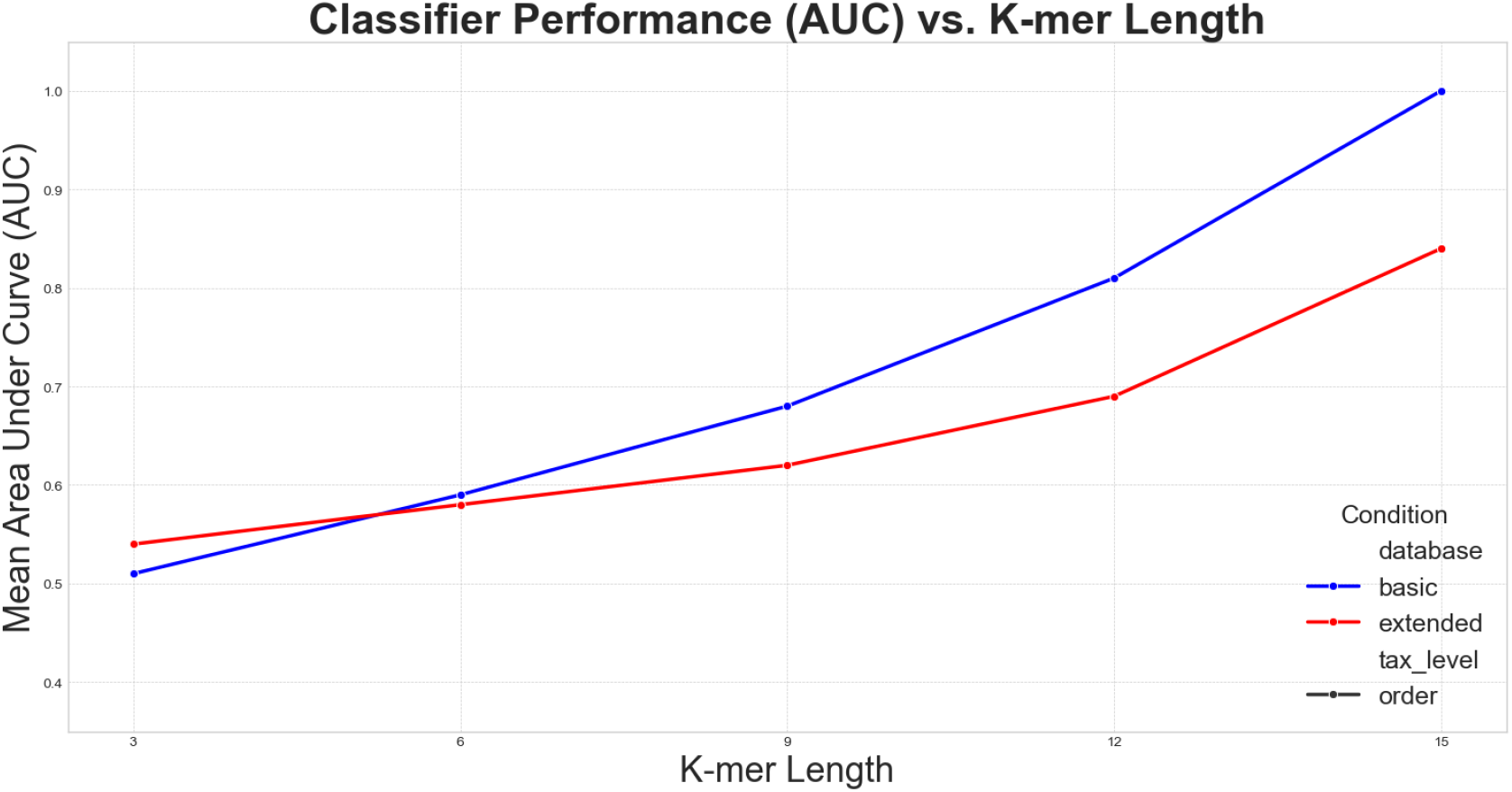
AUC against K-mer length (Order level)

When comparing the two databases, the model trained on the basic database often yielded slightly higher AUCs than the model trained on the extended database. This suggests that while both models improve with longer k-mers, the classification task itself becomes more challenging with a denser, more complex reference set. With many more closely related species in the extended database, the potential for ambiguity increases, leading to a slightly lower, but likely more realistic, measure of classification performance.

### Novelty thresholds across Database Scales

After determining the optimal log-probability for each model using Youden’s J statistic, we compared the values derived from the basic database against those from the extended database. Despite the vast difference in size and taxonomic complexity between the two databases, the calculated novelty thresholds were remarkably stable (Fig 2). For any given k-mer length and taxonomic level, the threshold value remained nearly identical regardless of which database was used for training. This finding suggests that the novelty thresholds (at various taxonomic levels) do not significantly change with database depth – as long as a taxonomic lineage has representation, the threshold value for determining that it is “in-distribution” remains fairly fixed.

**Figure 2.**
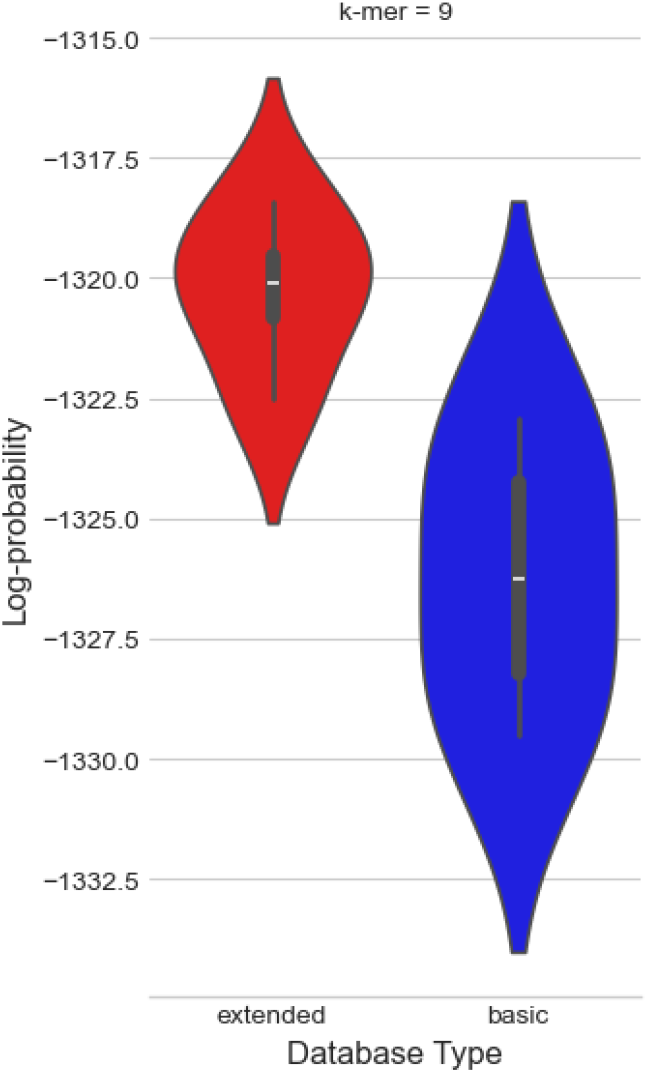
Optimal thresholds across taxonomic levels (9-mers)

### Application to a Human Gut Metagenome

To assess the practical performance of our framework, we applied the 9-mer models from both the basic and extended databases to classify reads from a real human gut metagenome.

Interestingly, the model trained on the smaller basic database classified a higher proportion of reads as “known” (98.09%) compared to the extended model (92.77%). To confirm this finding was driven by the models’ intrinsic properties and not minor variations in their thresholds, a sensitivity analysis was performed. Applying a single, common threshold to both models’ output consistently resulted in the basic model classifying more reads as “known”. (Table 1)

**Table 1:** Proportion of reads classified as “known” under different thresholds and models.

| Threshold Used | Model Used | % Known |
| --- | --- | --- |
| -1049.62 (extended phylum) | Extended | 92.77 |
| -1049.62 (extended phylum) | Basic | 96.41 |
| -1053.81 (basic phylum) | Extended | 95.35 |
| -1053.81 (basic phylum) | Basic | 98.09 |

To better understand the mechanism behind this disparity, we found that 75% of the reads classified by the extended model produced a more negative log-probability score than the basic model, indicating a lower classification confidence in its predictions (Fig. 3).

**Figure 3.**
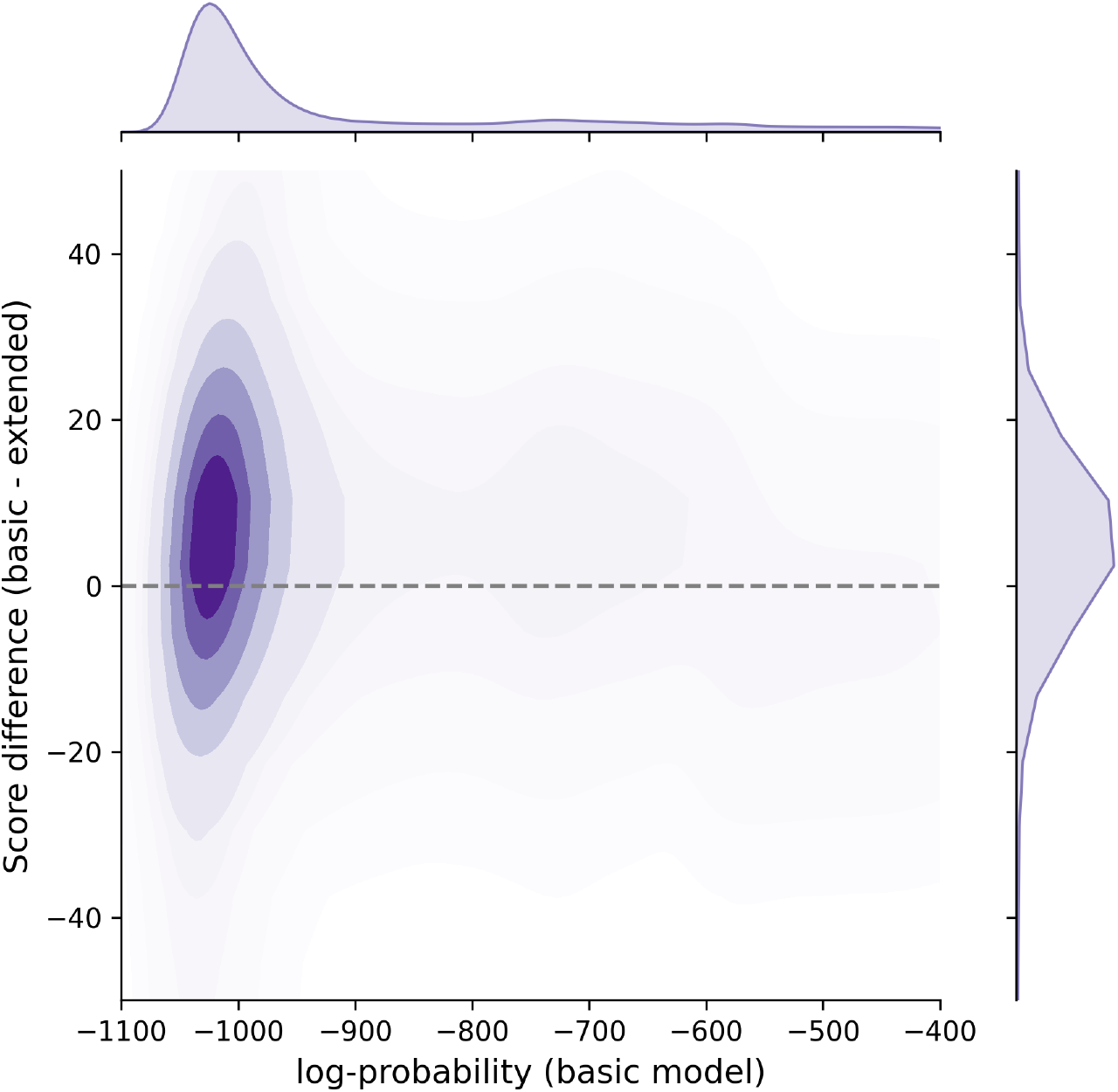
Score difference between the two 9-mer models

Taken together, these results provide strong evidence that the extended model operates with a higher standard of certainty. This higher standard is a direct result of its training on a denser taxonomic space, leading to a more conservative and likely more accurate estimate of the sample’s “known” fraction.

### Determining Optimal Thresholds

To pinpoint the most effective log-probability score for identifying novel sequences, we performed a Receiver Operating Characteristic (ROC) analysis on each model’s output. We generated an ROC curve using the scores from our test set, treating reads from the training set as the “known” positive class and reads from the holdout set as the “unknown” negative class. The optimal threshold was then identified as the point on this curve that maximized the **Youden’s J statistic**, calculated as *J* = sensitivity + specificity − 1. This process was repeated for each of the five independent trials for a given experimental condition (e.g., order level with 12-mers Fig 4). The final, definitive threshold reported for that condition was then taken from the single trial that yielded the highest Youden’s J statistic, representing the best classification performance observed across the replicates.

**Figure 4.**
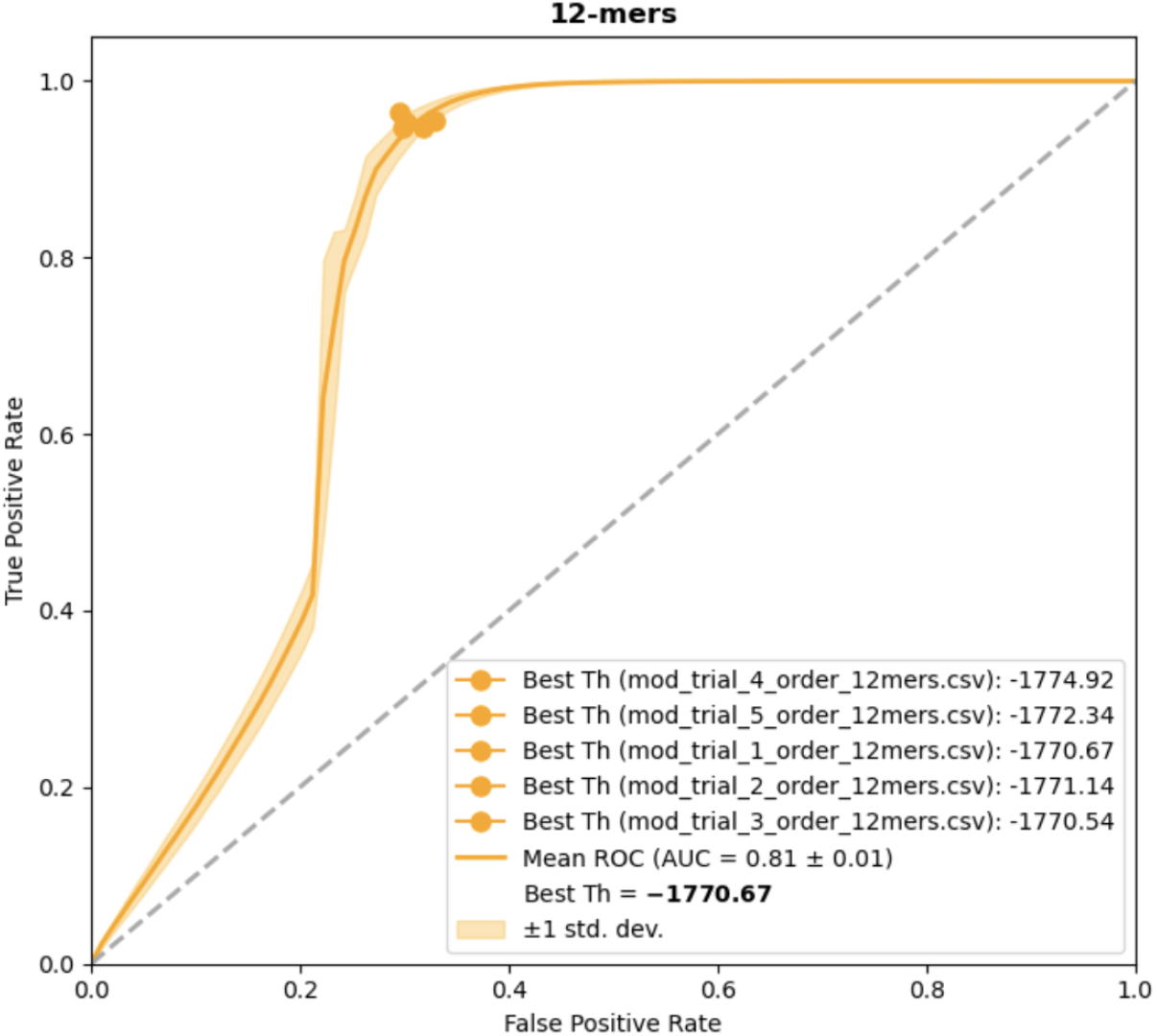
Order level 12-mer ROC

## Abbreviations

NBC: Naïve Bayes Classifier
NBC++: “Incrementalized” version of the Naïve Bayes Classifier

